# Autolysis affects the iron cargo of ferritins in neurons and glial cells at different rates in the human brain

**DOI:** 10.1101/2022.01.26.477869

**Authors:** Sowmya Sunkara, Snježana Radulović, Saška Lipovšek, Christoph Birkl, Stefan Eggenreich, Anna Maria Birkl-Toeglhofer, Maximilian Schinagl, Daniel Funk, Michael Stöger-Pollach, Johannes Haybaeck, Walter Goessler, Stefan Ropele, Gerd Leitinger

**Affiliations:** Gottfried Schatz Research Center, Division of Cell Biology, Histology and Embryology, Research Unit Electron Microscopic Techniques, Medical University of Graz, Graz, Austria; Faculty of Medicine, University of Maribor, Maribor, Slovenia; Department of Biology, Faculty of Natural Sciences and Mathematics, University of Maribor, Maribor, Slovenia; Faculty of Chemistry and Chemical Engineering, University of Maribor, Maribor, Slovenia; University Clinic for Neuroradiology, Medical University of Innsbruck, Innsbruck, Austria; Neuroimaging Research Unit, Department of Neurology, Medical University of Graz, Austria; Institute for Pathology, Neuropathology and Molecular Pathology, Medical University of Innsbruck, Innsbruck, Austria; Institute of Chemical Technologies and Analytics, Technische Universität Wien, Vienna, Austria; University Service Centre for Transmission Electron Microscopy (USTEM), Technische Universität Wien, Vienna, Austria; Diagnostic & Research Center for Molecular BioMedicine, Institute of Pathology, Medical University of Graz, Graz, Austria; Institute for Chemistry, University of Graz, Graz

**Keywords:** Ferritin, human brain, energy-filtered transmission electron microscopy, quantitative magnetic resonance imaging, inductively coupled plasma mass spectrometry, frontal gray matter, frontal white matter, putamen, globus pallidus, autolysis, *post-mortem*

## Abstract

Iron is known to accumulate in neurological disorders, so a careful balance of the iron concentration is essential for healthy brain functioning. An imbalance in iron homeostasis could arise due to the dysfunction of the proteins involved in iron homeostasis. Here, we focus on ferritin – the primary iron storage protein of the brain. Though it is known that glial cells and neurons differ in their concentration of ferritin, the change in the number of iron-filled ferritin cores or their distribution between different cell types during autolysis has not been revealed yet. Here, we show the cellular and region-wide distribution of ferritin in the human brain using state-of-the-art analytical electron microscopy. We validated the concentration of iron-filled ferritin cores to the absolute iron concentration measured by quantitative MRI and inductively coupled plasma mass spectrometry. We show that ferritins lost iron from their cores with progressing autolysis whereas the overall iron concentrations were unaffected. Though the highest concentration of ferritins was found in glial cells, we found that as the total ferritin concentration increased in a patient, ferritin accumulated more in neurons than in glial cells. Collectively our findings point out the unique behaviour of neurons in storing iron during autolysis and explain the differences between the absolute iron concentrations and iron-filled ferritin in a cell-type-dependent fashion in the human brain.

**Significance statement:** Balance of the iron load of the brain is crucial to preventing neurodegenerative disorders. Our study establishes a relation between autolysis, iron, and ferritin in the human brain with emphasis on the role of different cells in ferritin storage. We demonstrate that the iron load of ferritins does not correlate with mean iron concentrations during autolysis. Neurons retain more iron-loaded ferritin than glial cells with increasing ferritin count, which may make neurons more susceptible and exacerbate neuronal loss during iron overload. Neurons are also depleted of iron-loaded ferritin cores faster than glial cells during autolysis, demonstrating their unique role in iron storage. This paves the way to understanding the respective roles of neurons and glial cells in preventing or promoting neurodegeneration.

## Introduction

Non-heme iron accumulates in the normal ageing brain until the fourth decade (Hallgren and Sourander, 1958). The rate of accumulation and the absolute iron concentration vary strongly across brain regions with the highest concentrations in the basal ganglia, including the putamen and the globus pallidus (GP) (Hallgren and Sourander, 1958; Krebs et al., 2014). The cause of iron accumulation is still unclear, in particular since much more iron is stored than metabolically relevant.

Basal levels of iron are essential in an array of functions in the brain: e.g. myelin synthesis, as cofactors for enzymatic reactions (Codazzi et al., 2015), and synaptic plasticity (Muñoz et al., 2011). Labile iron ions act as free radicals, induce reactive oxygen species (Halliwell, 2006) and can lead to neuroferritinopathy followed by neurodegeneration (Friedman et al., 2011). In a normal functioning brain, excess iron that is unused is stored in its primary storage protein, ferritin (Wang and Pantopoulos, 2011; (Finazzi and Arosio, 2014)), shielding the brain from the potential iron toxicity.

Ferritin is at the heart of the iron regulation. Ferritin acts as a buffer that stores and releases iron upon cellular demand, which is crucial in the prevention of ferroptosis (apoptosis induced by iron overload) (Jiang et al., 2021) and other detrimental effects caused by free iron. Ferritin contains between 500 to 5000 ferric iron atoms in its inner core (Harrison and Arosio, 1996; Iancu, 2011; Jian et al., 2016). The size of the ferritin core in electron micrographs ranges from 5.3 to 11 nm (Iancu, 2011) depending on various factors such as iron load, contrasting agent, hydration, or the focus of the micrograph. Despite the difference in the concentrations of total iron and ferritin-bound iron between brain areas, the size of ferritin cores remains largely uniform (Friedman et al., 2011).

Ferritin takes a unique position amongst the iron-binding proteins, owing to its size and cargo capacity. Other iron-containing proteins like transferrin and haemoglobin contain only 2 or 4 iron atoms respectively (Giometto et al., 1993; Wagner et al., 2003; Marengo-Rowe, 2006), and hence are below the detection threshold for energy-filtered transmission electron microscopy (EFTEM), our method of choice for ferritin detection. Conversely, hemosiderin is composed of conglomerates of clumped ferritin particles, denatured proteins, and lipids with a size of only 1 - 2 μm (Wagner et al., 2003). In view of that, hemosiderin is easily distinguished by size from ferritin but is rarely detected in the normal brain.

*Post-mortem* studies have the advantage that tissue is available for a variety of measurements and procedures, including destructive ones. But autolysis sets off immediately after death, causing a breakdown of cells and tissues, changing structural components, and making it difficult to identify them (Sele et al., 2019), reviewed in (Lewis et al., 2019).

The total iron concentration in freeze-dried tissue samples can be determined precisely by inductively coupled plasma mass spectrometry (ICP-MS), e.g. in (Krebs et al., 2014). However, this method is destructive. A non-invasive and thus non-destructive way to measure spatial iron concentrations is quantitative Magnetic Resonance Imaging (qMRI). qMRI allows mapping the local transverse relaxation rate R_2_*, which has been validated as a measure for iron content in brain tissue using ICP-MS (Langkammer et al., 2010). However, neither method provides information on the amount of iron stored in ferritins or its cellular distribution. We thus established a workflow using EFTEM for localising and quantifying the iron–filled cores of ferritins. We combined the three methodological approaches in our study to obtain complementary insights into the effects of autolysis on iron storage within ferritin cores.

For this, we investigated the concentrations of iron and ferritin cores filled with iron in the human brain samples *post-mortem* at the region and cellular level. We observed that autolysis influences the loss of iron-filled ferritin cores in a time-dependent manner. Moreover, our study reveals different rates of loss of iron–filled cores between neurons and glial cells during autolysis.

## Materials and methods

### Human brain samples

The samples used here are collected from six deceased human subjects who had to undergo routine brain autopsy at the Diagnostic and Research Institute of Pathology of the University Hospital Graz (approved by the Ethics Committee of the Medical University of Graz, votum number 28-549 ex 15/16). The subjects had not suffered from a known neurological disorder. The sex of the subjects was not considered for this study. To ensure that the analyses were not confounded by age, the specimens were collected from patients with ages ranging from 61-86. At this age saturation in iron accumulation has already been reached (Hallgren and Sourander, 1958). The age and *post-mortem* intervals (PMI) are given in Table 1. One-half of the brain was preserved for quantitative magnetic resonance imaging (qMRI) while four brain areas were dissected out of the other half and further divided into samples for EFTEM and ICP-MS. The brain regions selected included two regions with the lowest iron concentrations - the frontal gray matter (FGM), the superficial frontal white matter (FWM), and two regions with higher concentrations of iron - the putamen, and the globus pallidus (GP) (Bilgic et al., 2012; Ward et al., 2014; Ramos et al., 2014). Once the tissue was cleared for research purposes by the pathologists, samples were dissected out and either immersed in fixative for EM, or placed into a plastic tube, weighed, and frozen for ICP-MS. The brain hemisphere for qMRI remained unfixed.

**Table 1.** Patient information

| Patient code | Age [years] | PMI [h] |
| --- | --- | --- |
| 1 | 73 | 06:30 |
| 2 | 72 | 9 |
| 3 | 61 | 16 |
| 4 | 69 | 18 |
| 5 | 86 | 20 |
| 6 | 81 | 24 |

### Ferritin Isolation

Ferritin was isolated from human *post-mortem* brain samples in a series of purification steps. In brief, the brain sample weighing approximately 10g immersed in 10mM Tris-HCL, 150mM NaCl containing protease inhibitor was homogenized using Turrax homogenizer IKA T −10. Followed by centrifugation at 10000g for 30-60 min at 4°C and sonication for 2 min at 4°C using a Branson Sonifier 250. The sample was centrifuged again at 10000 g for 30-60 min at 4°C retaining the supernatant for further purification. The supernatant was heated up to 70-75°C for 10 mins under constant stirring and immediately cooled. 75% saturated ammonium sulfate was used to precipitate the proteins in the supernatant overnight at 4 °C. The precipitated proteins were collected the following day by centrifugation at 10000 g for 60 min at 4 °C. This was followed by size exclusion chromatography and density gradient centrifugation. Isolated ferritin was loaded onto a carbon-coated nickel grid and stained with 1% uranyl acetate to negatively contrast the sample before visualization by EM.

### Electron Microscopy

Following the autopsy, 6-10 sub-samples of approximate wet weight 0.1-0.2 g (maximum dimensions 1×1×1 mm^3^) were cut out of the samples and immersed in chemical fixative containing 2% formaldehyde, 2% glutaraldehyde in 0.1 M sodium cacodylate buffer. After fixation, the samples were rinsed in the same buffer, postfixed in 1% osmium tetroxide solution in the same buffer, dehydrated in a graded series of alcohol, immersed in propylene oxide, and embedded in TAAB embedding resin (TAAB, Aldermaston, UK). After curing for 3 days at 60 °C, thin sections were cut using a Leica UC6 or a Leica UC7 ultramicrotome at a thickness of either 60 or 70 nm. The sections were contrasted with platinum blue and lead citrate solutions. One block was randomly selected from each brain area for imaging and elemental mapping.

EM was performed with a Thermo Fisher Tecnai G2 20 electron microscope operated at 200 keV. An Ametek Gatan Quantum GIF energy filter with Ametek Gatan Ultrascan 1000 XP camera was used for EFTEM, and a bottom-mounted Ametek Gatan Ultrascan 1000 camera for bright field imaging. For elemental mapping, three windows – Pre-edge 1, Pre-edge 2, and Post-edge – were made at 80000x magnification, a binning of 2, a slot width of 40 eV, and an exposure time of 30s. The energy losses were: 718 eV for the iron L – post-edge window, and 633 eV and 673 eV for the two pre-edge windows. Drift correction was done manually, and both an iron L – elemental map and an iron L – jump-ratio were computed on Digital Micrograph (Gatan, Inc) using respectively the three-window method and the two-window method. The iron cores inside the ferritins are visible as bright spots using EFTEM.

An unbiased sampling protocol described earlier (; Wernitznig et al., 2019) automatically selects 60 random locations on each sample determined using a random number generator. The elemental maps were accompanied by overview micrographs taken from the same locations using the bright field mode at 3500x magnification.

### Ferritin quantification and cell type assessment

Four independent researchers (SS, SnR, SL, and GL) each quantified the number of iron-loaded ferritins and assessed the cell types. Ferritins were only counted if they were clearly visible as a bright spot on both the iron L – jump-ratio and the iron L – elemental maps and if at least three of the four researchers agreed on their number. In case of disagreement, the images were analysed again simultaneously. If the number of ferritins exceeded 20 in any map, the mean value of the counts from four researchers was considered as the final count. Cell types were identified on the bright-field images and only noted if three of the four researchers agreed. For cell identification, the criteria described by (Nahirney and Tremblay, 2021) and the Boston University web atlas https://www.bu.edu/agingbrain/ (download 2022-01-14) were used. A total of 1440 iron L – elemental maps and 1440 iron L – jump ratios were generated and used for this study.

### Quantitative Magnetic Resonance Imaging

MR imaging of the extracted brain hemisphere was performed using a 3.0 Tesla scanner (MAGNETOM PRISMA, Siemens Healthineers, Erlangen, Germany). A phased array head coil with 20 elements was used. For acquiring R_2_* relaxation data, a two-dimensional radiofrequency-spoiled multi-echo gradient-echo sequence with 6 equally spaced echoes was applied (repetition time = 1150 ms, first echo time = 4.92 ms, echo spacing = 5.42 ms, flip angle = 15°; field of view = 256 x 256 mm^2^; in-plane resolution = 0.75 x 0.75 mm^2^; 2.4-mm-thick sections covering the entire hemisphere). Generalized auto-calibrating partially parallel acquisition (GRAPPA) was performed with an acceleration factor of 2.

To reduce phase dispersion effects and to increase B0 homogeneity, second-order shimming was applied. R_2_* maps were calculated with the relaxometry toolbox(2016).

### Inductively Coupled Plasma Mass spectrometry

All solutions were prepared with ultrapure water (18.2 MΩ*cm, Merck Millipore, Bedford, USA). Nitric acid (>65 % p.a.) was further purified via sub-boiling. For quantification, single elements standards (ROTI^®^Star, Carl Roth GmbH + Co. KG, Karlsruhe, Germany) were used.

Brain tissue was freeze-dried to constant mass with a Gamma 1-16 LSC freeze-dryer (Martin Christ GmbH, Osterode am Harz, Germany) and the wet/dry mass ratio was determined. Freeze-dried samples were weighed to 0.1 mg into the 12 mL quartz vessels of a microwave-heated autoclave, UltraCLAVE IV (EMLS, Leutkirch, Germany). After the addition of 3 mL subboiled nitric acid and 2 mL of water the quartz vessels were placed in the 40 positions rack of the autoclave and the autoclave was pressurized with Argon to a pressure of 4*10^6^ Pa. The following microwave heating program was applied: from room temperature to 80°C in 5 minutes, from 80°C to 150°C in 15 minutes, from 150°C to 250°C in 20 minutes, and 250°C for 30 minutes. After cooling to 80°C, the pressure was released and the samples were transferred to 50 mL polypropylene tubes (CELLSTAR^®^, blue screw cap, Greiner Bio-One GmbH, Kremsmuenster, Austria) and filled to the mark.

For the quantification of iron (Fecalibration curves were prepared from 0.0100 to 10.0 (Mg, Ca, Fe). Germanium (Ge) at a concentration of 200 μg/L was added as an internal standard online before the nebulizer. The samples were pumped through a peristaltic pump tubing with an i.d. of 1.05 mm and the internal standard with a tubing of 0.19 mm. The elemental iron was quantified with an inductively coupled plasma mass spectrometer (ICP-MS, Agilent 7700x, Agilent Technologies, Waldbronn, Germany) at a mass-to-charge ratio of Fe m/z = 56. All isotopes were measured in the collision mode using He at a flow rate of 5.0 ml/min. The determined concentrations were calculated for fresh tissue.

The accuracy of the determined concentrations was confirmed with the certified material bovine muscle (RM 8414; NIST; now BOVM-1 NRCC).

### Statistical Analysis

Statistical analysis was performed using GraphPad Prism 9.2.0. One-way ANOVA with a Tukey’s multiple comparisons test was used for statistical analysis with * p < 0.05; *** p < 0.001; **** p < 0.0001 for figures 2,3,5 and 8. For all comparison plots of data from three methodological approaches in figures 6, 7, and 9, a simple linear correlation analysis was performed.

## Results

### EFTEM elemental mapping allows visualizing the ferritins’ cores in the human brain

Ferritins consist of a shell and an iron-filled core, and the core is visible as an electron-dense body in negative contrast electron micrographs of isolated ferritins (Fig. 1B). The same core is visible as a bright spot in iron L – elemental maps (Fig. 1C).

**Figure 1.**
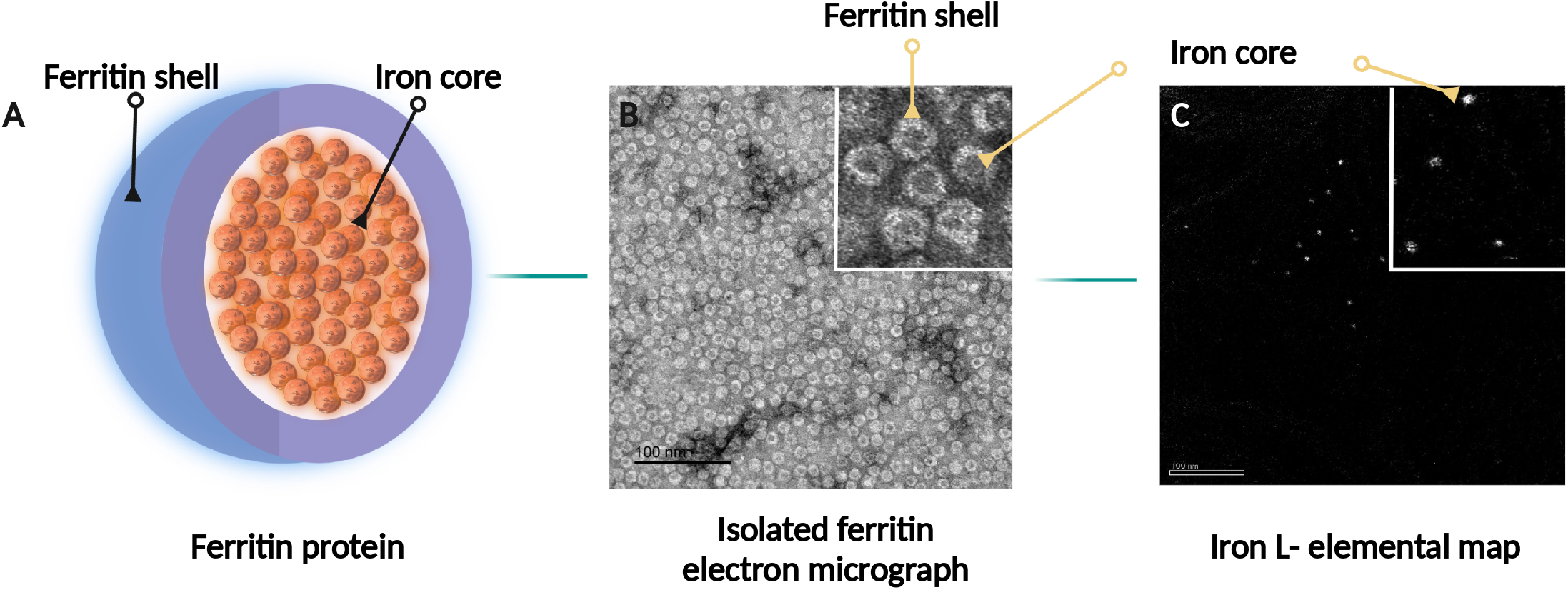
Visualization of the iron-core of ferritin. **A,** Schematic diagram of the molecular architecture of ferritin. **B**, Negative contrasted electron micrograph of isolated ferritin from the human brain. The ferritin shell appears as white rings shielding the electron-dense iron core (inset). **C,** Iron L – elemental map displaying the iron cores as bright spots.

To visualize the ferritin cores in electron micrographs, we generated iron L – elemental maps and iron L – jump-ratios of brain samples for four different brain areas in each of six different deceased subjects. The maps consistently exhibited ferritins as bright spots of 5-10 nm in diameter in all four examined brain regions (Fig. 2A-D). The size of the spots corresponded to the published sizes of ferritin cores (Iancu, 2011), indicating that iron-filled ferritin cores could be visualized with EFTEM.

**Figure 2.**
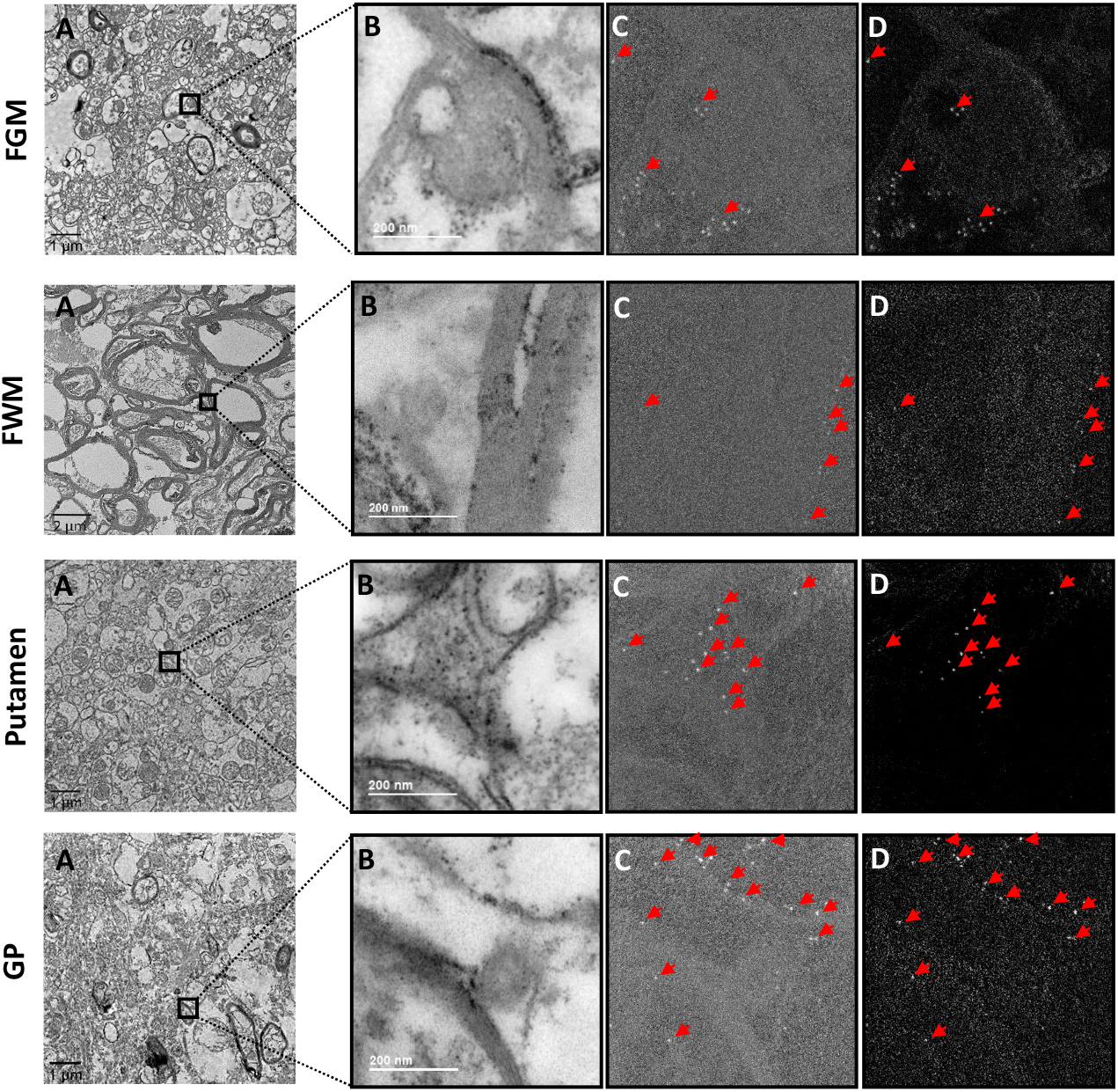
EFTEM mapping of iron in different regions in post-mortem human brain sample. **A**, Bright-field electron micrographs of overview images of the FGM region, the FWM region, the putamen region, and the GP region. **B**, Inverted post-edge micrographs. **C**, iron L – jump-ratios. **D**, corresponding iron L – elemental maps of the same region as B. Ferritin filled with iron core appear as bright spots. Examples are marked by red arrows. Corresponding iron L – jump-ratios and iron L – elemental maps (C, D) show ferritin present in clusters in the cells.

### Ferritin cores were clustered in distinct cells

We next counted the number of ferritin cores in samples taken from FGM, FWM, putamen, and GP of each patient. The counts were made within 60 micrographs of 556 x 556 nm^2^ dimensions and showed that the ferritins were not evenly distributed but clustered in certain micrographs (Fig. 3). We observed that the majority of ferritin accumulated as clusters as shown in figure 1, inside the cells. At a closer examination, it became apparent that certain cells accumulated ferritins, whereas others, often neighbouring cells, did not accumulate ferritins (Fig. 2). None of the ferritins appeared to be extracellular. We were able to assess the cell type (neuron, oligodendrocyte, astrocyte, or other unidentified glial cells) in 2,195 (62 %) of the 3,537 ferritin cores that we had identified in five out of six samples. 471 additional ferritin cores were identified in the sixth sample with a *post-mortem* interval (PMI) of 24 h, but in this sample, autolysis had progressed considerably, preventing us from faithfully identifying cell types. Accordingly, this sample was excluded from cell type identification experiments.

**Figure 3.**
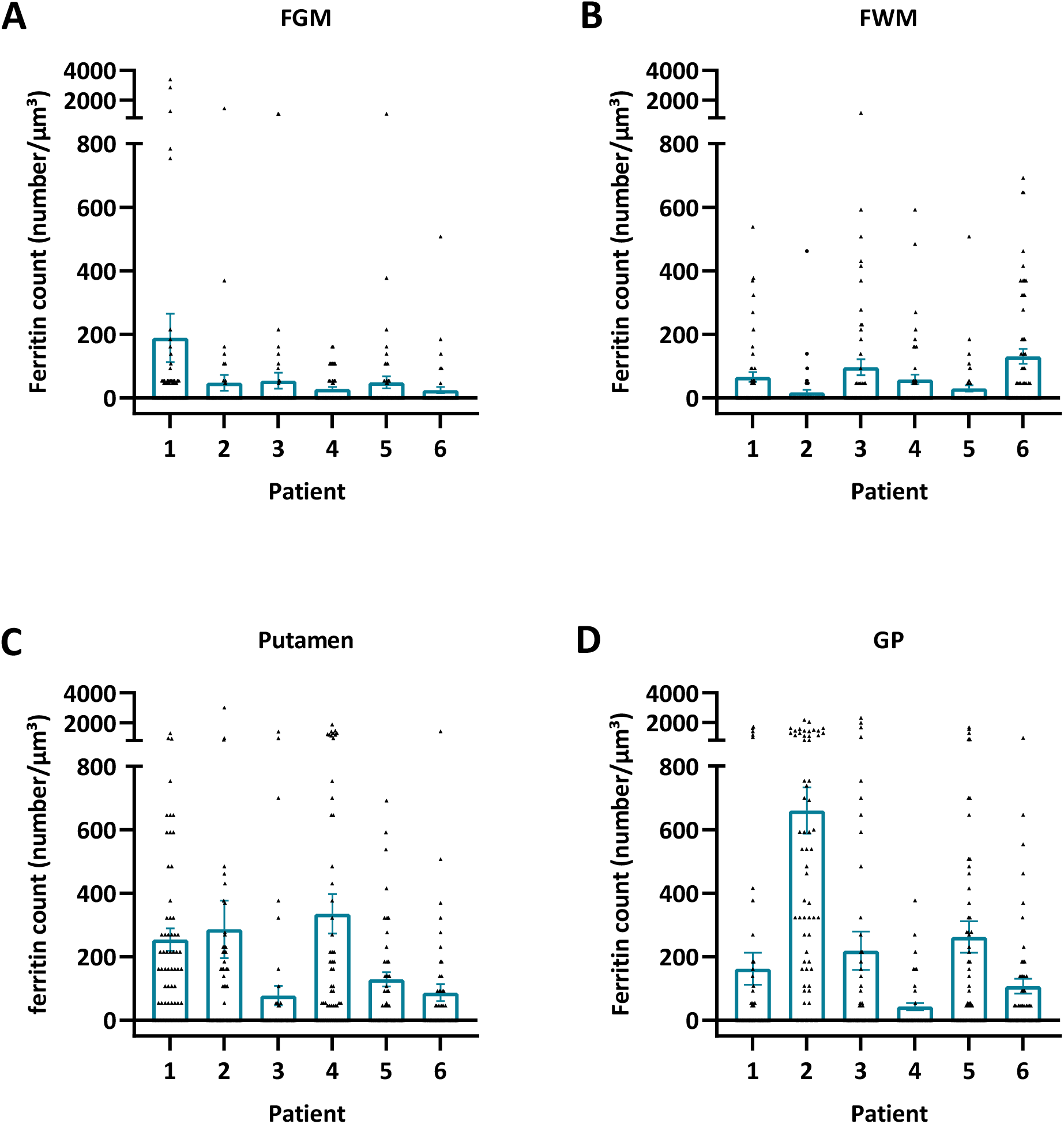
The ferritin concentration in different brain regions varies between patients 1-6. **A**, FGM. **B**, FWM. **C**, Putamen. **D**, GP. Data are shown as bar graphs with whiskers representing the mean ± SEM. Each dot represents the ferritin concentration measured in one iron L - elemental map, (n = 60 for each patient in each brain region). One-way ANOVA with Tukey’ s multiple comparisons test was used.

### Ferritins accumulate in the putamen and globus pallidus

We next calculated the number of ferritins per μm^3^ from the micrographs’ 3D dimensions. Plotting ferritin concentration in each brain region for each patient revealed that either the putamen, GP, or both, had a significantly higher concentration of ferritins than either the FGM or FWM in five out of six patients (Fig. 4). Further, between the putamen and GP (4 out of 6 patients) significant differences were found (Fig. 4). The GP had a significantly higher concentration compared to the former in three out of six patients and the putamen had a significantly higher ferritin concentration than the GP in one out of six patients (Fig. 4).

**Figure 4.**
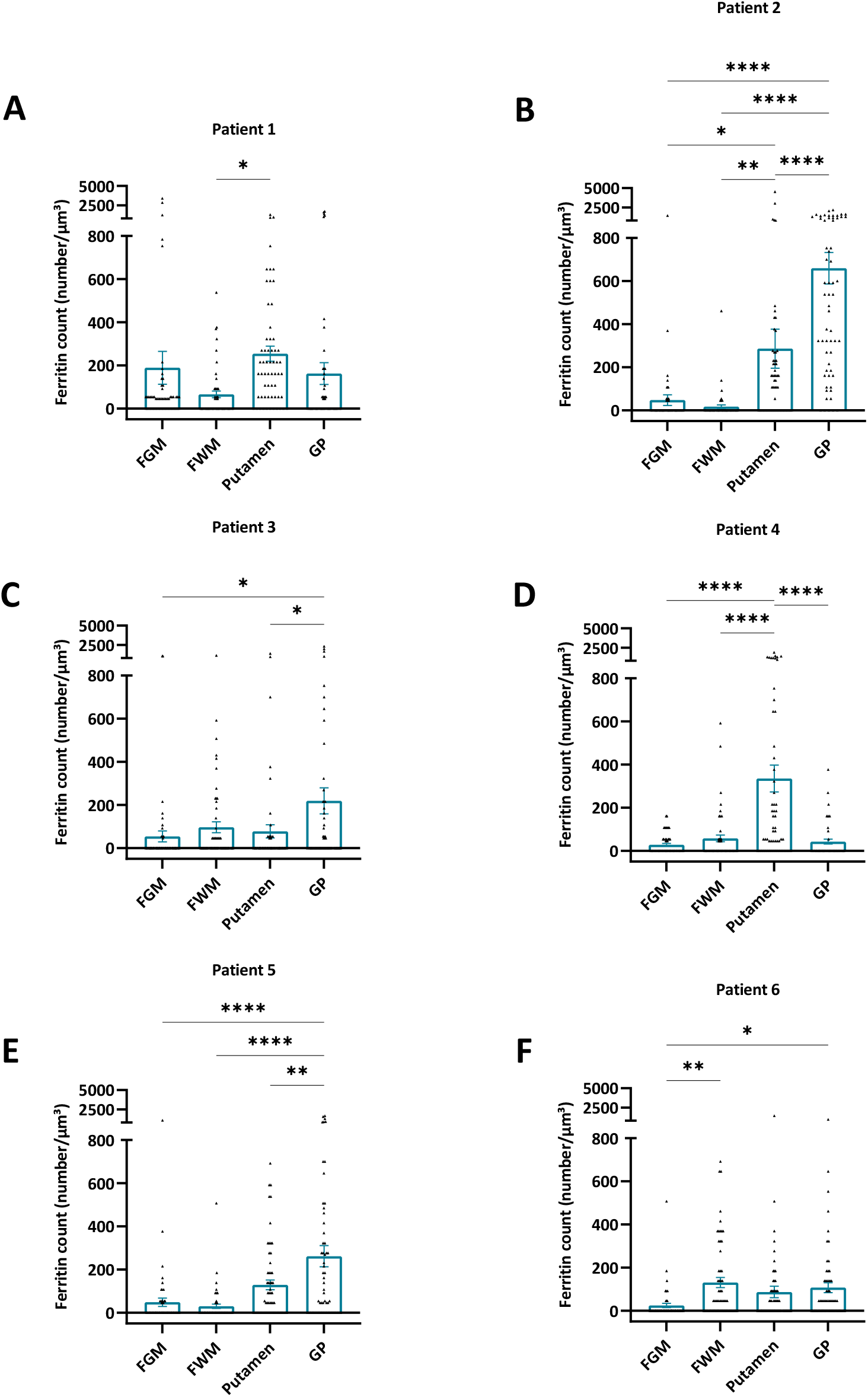
The ferritin concentration is often, but not always, higher in the basal ganglia than in the frontal gray or frontal white matter. Graphs show the ferritin concentration in the FGM, FWM, putamen, and GP in **A**, patient 1, **B**, patient 2, **C**, patient 3, **D**, patient 4, **E**, patient 5, and **F**, patient 6. Data are shown as bar graphs with whiskers representing the mean ± SEM. Each dot represents the ferritin concentration measured in one iron L - elemental map, respectively (n = 60 for each brain region and each patient). * p < 0.05; ** p < 0.01; **** p < 0.0001, One-way ANOVA with a Tukey’s multiple comparisons test.

### qMRI and ICP-MS data are on par with the EFTEM data

The ferritin concentrations counted from elemental maps were validated with the mean iron concentrations. The latter was determined by both mass spectrometry in adjacent tissue samples and by the proton transverse relaxation rate R^2^* – a qMRI measure proportional to the iron concentration – measured in the contralateral hemisphere (Fig. 5). Each of the three methodological approaches provided higher mean values (of all the six samples) in the putamen and GP than in FGM and FWM (Fig. 6A–C)). qMRI revealed a significant R_2_* increase in putamen and GP compared to FGM and FWM (Fig. 6B). Significant differences were found in the iron concentrations between the putamen and FGM using mass spectrometry (Fig. 6C).

**Figure 5.**
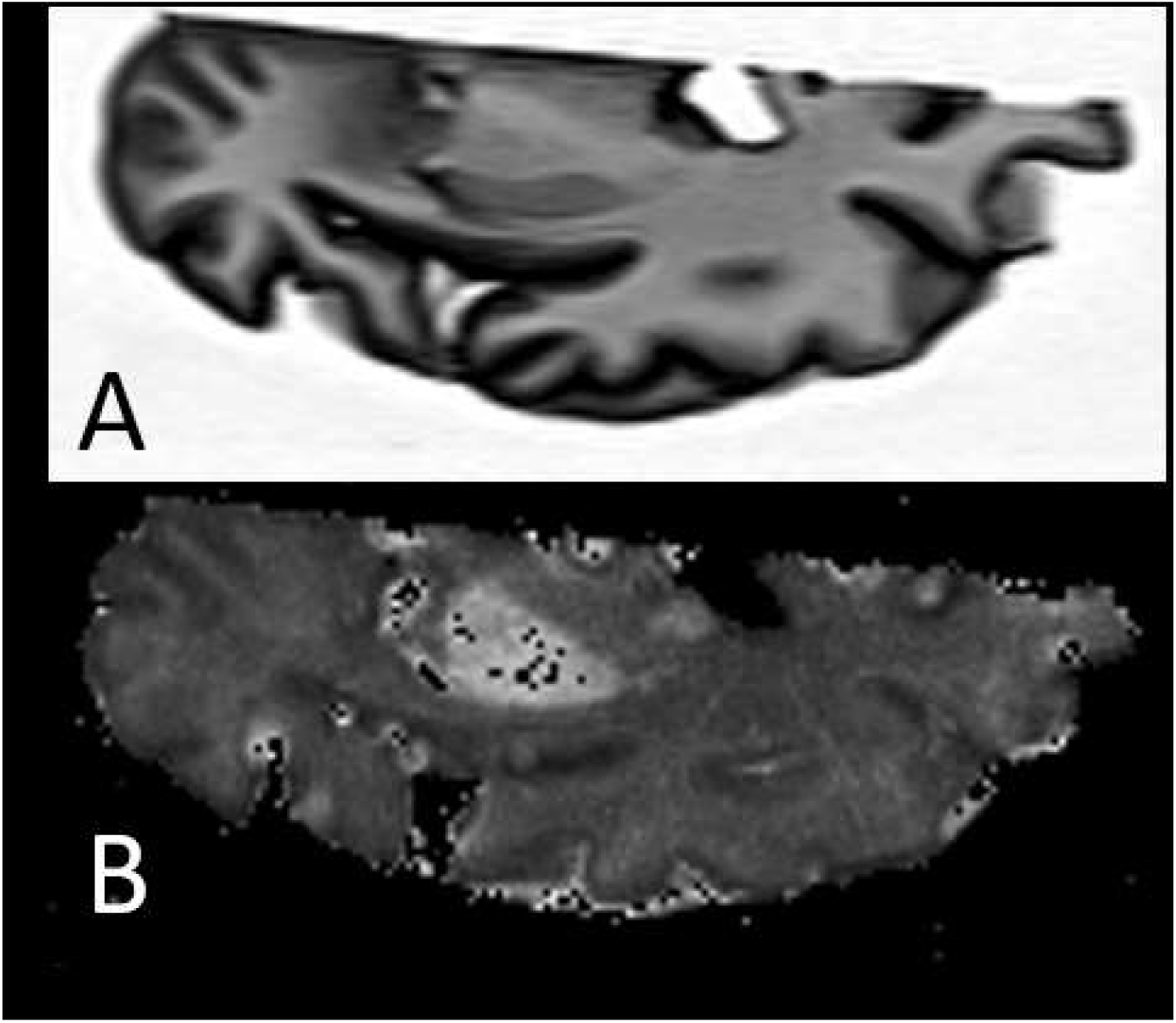
MRI scan showing an axial T1-weighted image of **A**, one cerebral hemisphere, and **B**, the corresponding R_2_* map. The bright areas in (B) are the putamen and the globus pallidus with higher R_2_* rates as a consequence of age- related iron accumulation.

**Figure 6.**
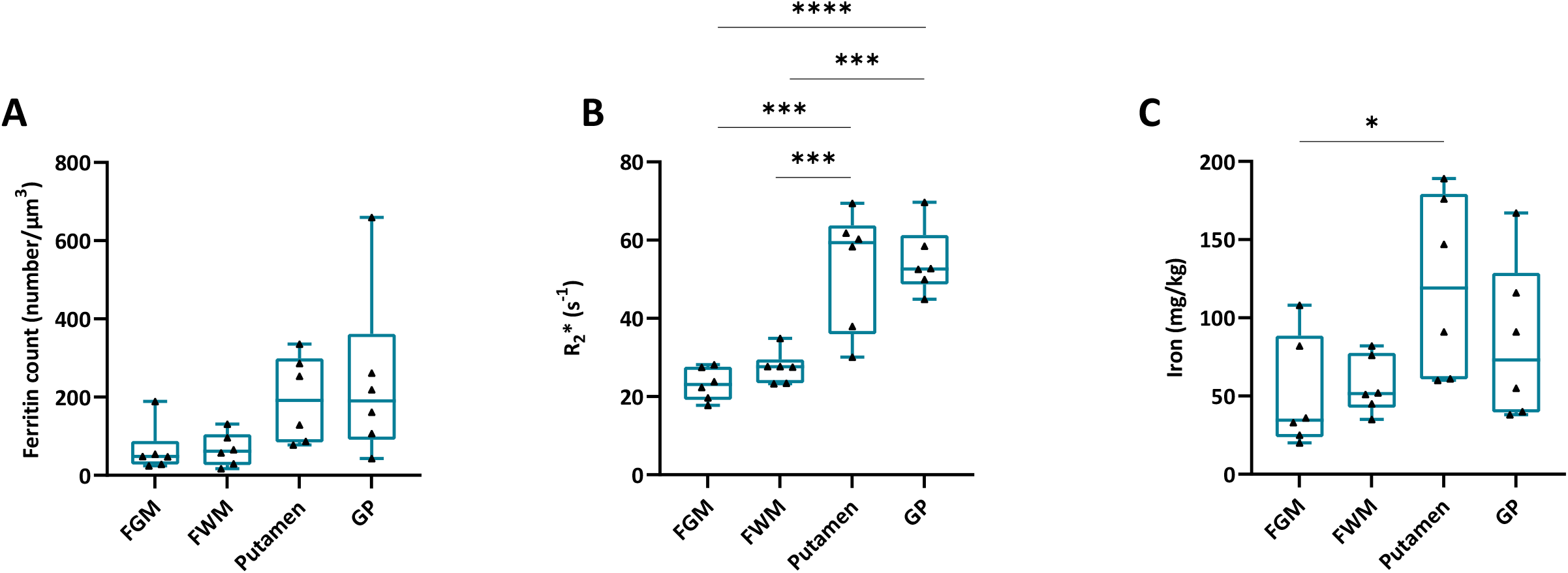
The differences in ferritin concentration determined by EFTEM between brain areas follow the same trend as the sample from the counter brain half in qMRI data and adjacent samples in ICP-MS data. **A**, Ferritin concentration in the FGM, FWM, putamen, and GP measured by EFTEM. **B**, Iron concentration in the FGM, FWM, putamen, and GP measured by ICP-MS. **C**, Proton transverse relaxation rate (R_2_*) in the FGM, FWM, putamen, and GP measured by qMRI. Data are shown as bar graphs with whiskers representing the mean ± SEM. Each dot represents the mean ferritin concentration measured in one patient, respectively (n = 6). * p < 0.05; ** p < 0.01; **** p < 0.0001, One-way ANOVA with a Tukey’s multiple comparisons test.

To compare the three different methodological approaches, the mean values of the measured brain regions from all six patients were correlated with each other. There was a high correlation between qMRI and ICP-MS results, each measuring the absolute iron concentration. The higher the R_2_* value, the higher the iron concentration (Fig. 7A, slope differs significantly from 0). However, we observed that the mean ferritin count obtained using EFTEM does not correlate with the iron concentration obtained with either qMRI (Fig. 7B) or ICP-MS (Fig. 7C). This apparent discrepancy prompted us to check whether the autolysis had an influence on the ferritin count or its cellular distribution.

**Figure 7.**
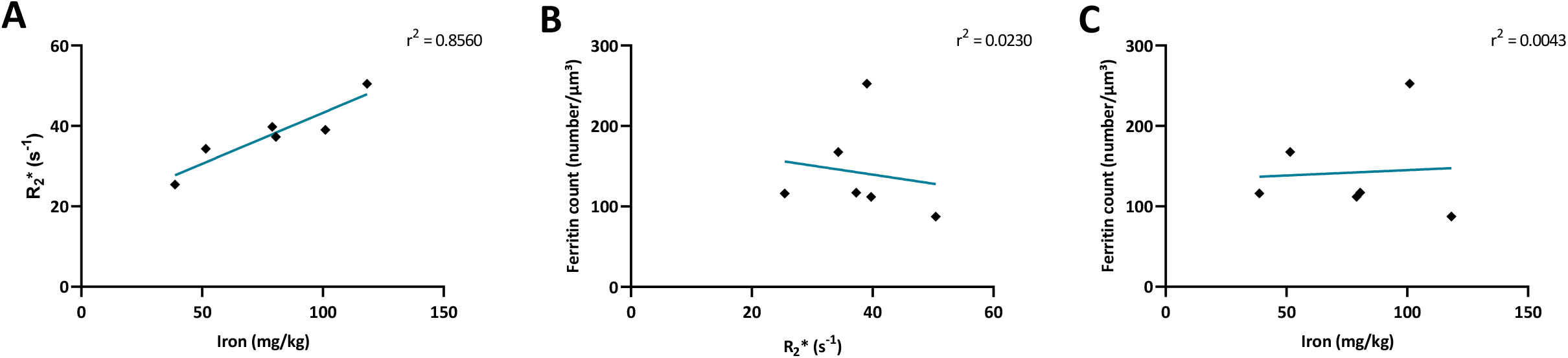
Correlation between mean concentrations of ferritin and iron measured via EFTEM, qMRI, and ICP-MS respectively. **A**, Proton transverse relaxation rate (qMRI) plotted over the iron concentration (ICP-MS) for each patient. The mean proton transverse relaxation rate of each patient correlates significantly with the mean iron concentration, r^2^ = 0.8560, p = 0.00082. **B**, Mean ferritin concentration (EFTEM) plotted over the proton transverse relaxation rate (R_2_*) (qMRI). **C**, Mean ferritin concentration (EFTEM) plotted over the iron concentration (ICP-MS). In contrast to (A), the mean concentration of ferritin particles does not correlate with proton relaxation rate (B), r^2^ = 0.0230, p = 0.7742 nor with the iron concentration (C), r^2^ = 0.0043, p= 0.9015. Each dot represents the mean ferritin concentration or the mean iron concentration measured in one patient, respectively (n = 6). All the plots are based on mean values of the FGM, FWM, the putamen, and the GP. * p < 0.05; ** p < 0.01; **** p < 0.0001, Linear correlation analysis.

### Autolysis enhances the unloading of the iron cargo from ferritin cores

We next verified whether putative autolysis that progresses with increasing PMI influenced the iron concentration, the storage of iron in ferritins’ core, or the cellular distribution of the ferritins.

Our analysis showed that the mean ferritin count was inversely correlated to the PMI (Fig. 8A). In contrast, neither the R_2_* value (Fig. 8B) nor the ICP-MS determined iron concentration correlated with the PMI (Fig. 8C). In other words, ferritin loses iron from its core with progressing autolysis while the absolute iron concentration remains stable in each patient. This indicates that the longer the *post-mortem* time, the more the ongoing autolysis progresses causing the unloading of iron from its storage protein.

**Figure 8.**
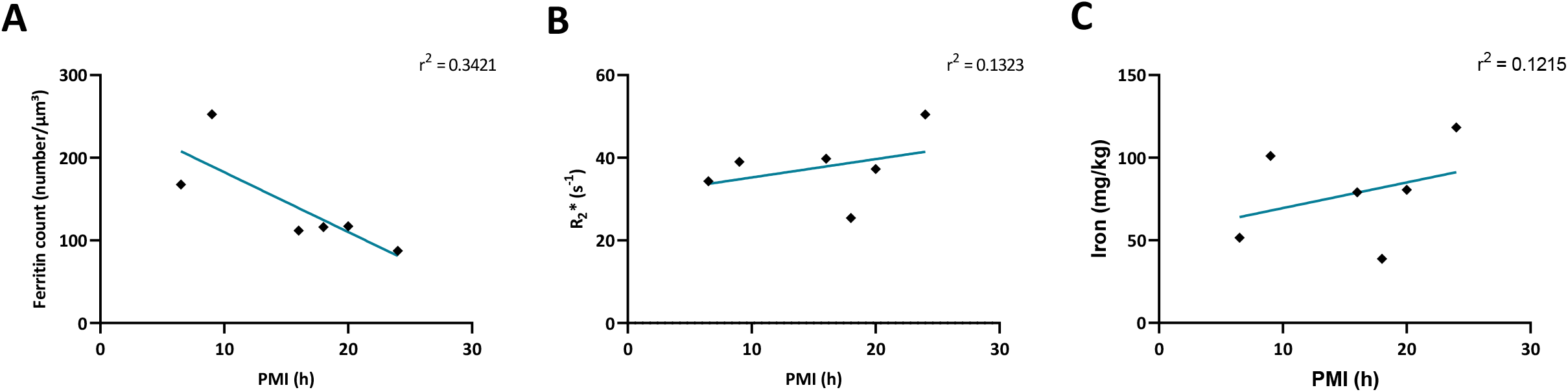
Ferritins disintegrate and their concentration is reduced over post-mortem time. **A**, Mean values of the ferritin concentration (EFTEM) in all the brain regions plotted against PMI, r^2^ = 0.6421, p = 0.0553. **B**, Mean values of the proton transverse relaxation rate (qMRI) in all the brain regions plotted against PMI, r^2^ = 0.1323, p = 0.4785. **C**, Mean values of the iron concentration (ICP-MS) in all the brain regions plotted against PMI, r^2^ = 0.1215, p = 0.4983. The ferritin concentration decreases with PMI in regions of the brain (A) whereas the total iron load in the four regions of the brain does not seem to be influenced by the PMI (B, C). Each dot represents the mean ferritin concentration or the mean iron concentration measured in one patient, respectively (n = 6). All the plots are based on mean values of the FGM, FWM, the putamen, and the GP. * p < 0.05, Linear correlation analysis.

### Cell type identification reveals that ferritins are highly concentrated in glial cells, especially oligodendrocytes

The counted ferritin was assigned to different cell types by four independent researchers. The cellular ferritin distribution showed that most of the 2195 ferritin cores that could be assigned to a specific cell type were found within glial cells. Oligodendrocytes represented the major cell type within this fraction (59 %). 17 % were found in neurons, 8% in astrocytes, and 1.7% in microglial cells (Fig. 9).

**Figure 9.**
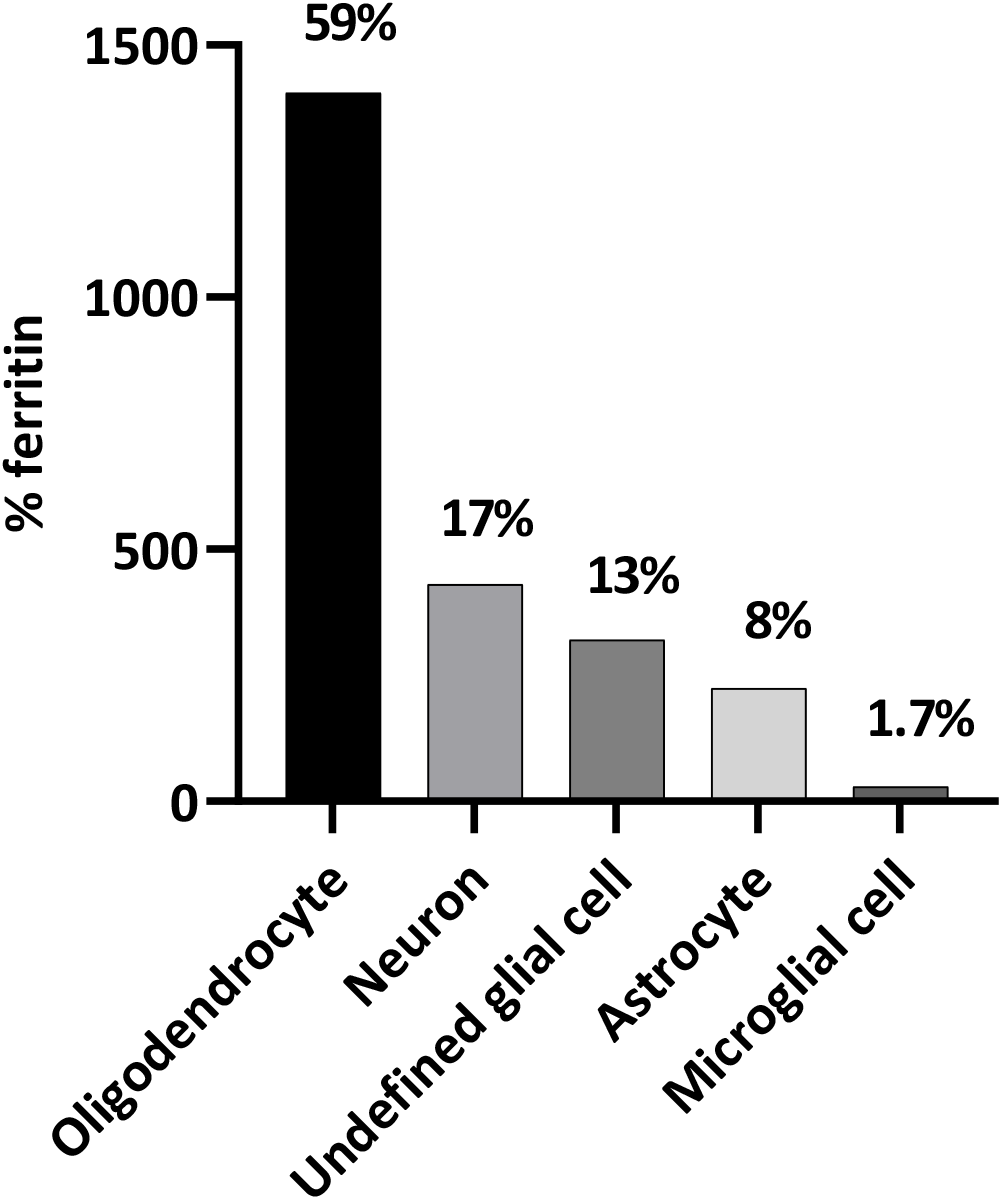
Ferritin is predominately stored in glial cells (82%) and especially in oligodendrocytes (59%); followed by 8% in astrocytes and 1.7% in microglial cells of the identified glial cells. 17 % is stored in neurons. The graph shows those ferritins in the FGM, FWM, putamen, and GP of patients 1-5 whose cell type could be determined. One-way ANOVA with Tukey’s multiple comparisons test was used for statistical analysis.

### Ferritin retention in neurons varies with both autolysis and the total ferritin count

Next, we studied the influence of autolysis on the iron load within ferritins in neurons. We found that the percentage of iron-filled ferritin cores found using EFTEM in neurons was inversely proportional to the PMI (Fig. 10A), indicating that unloading of ferritins during autolysis is faster in neurons than in glial cells (Fig. 10B).

**Figure 10.**
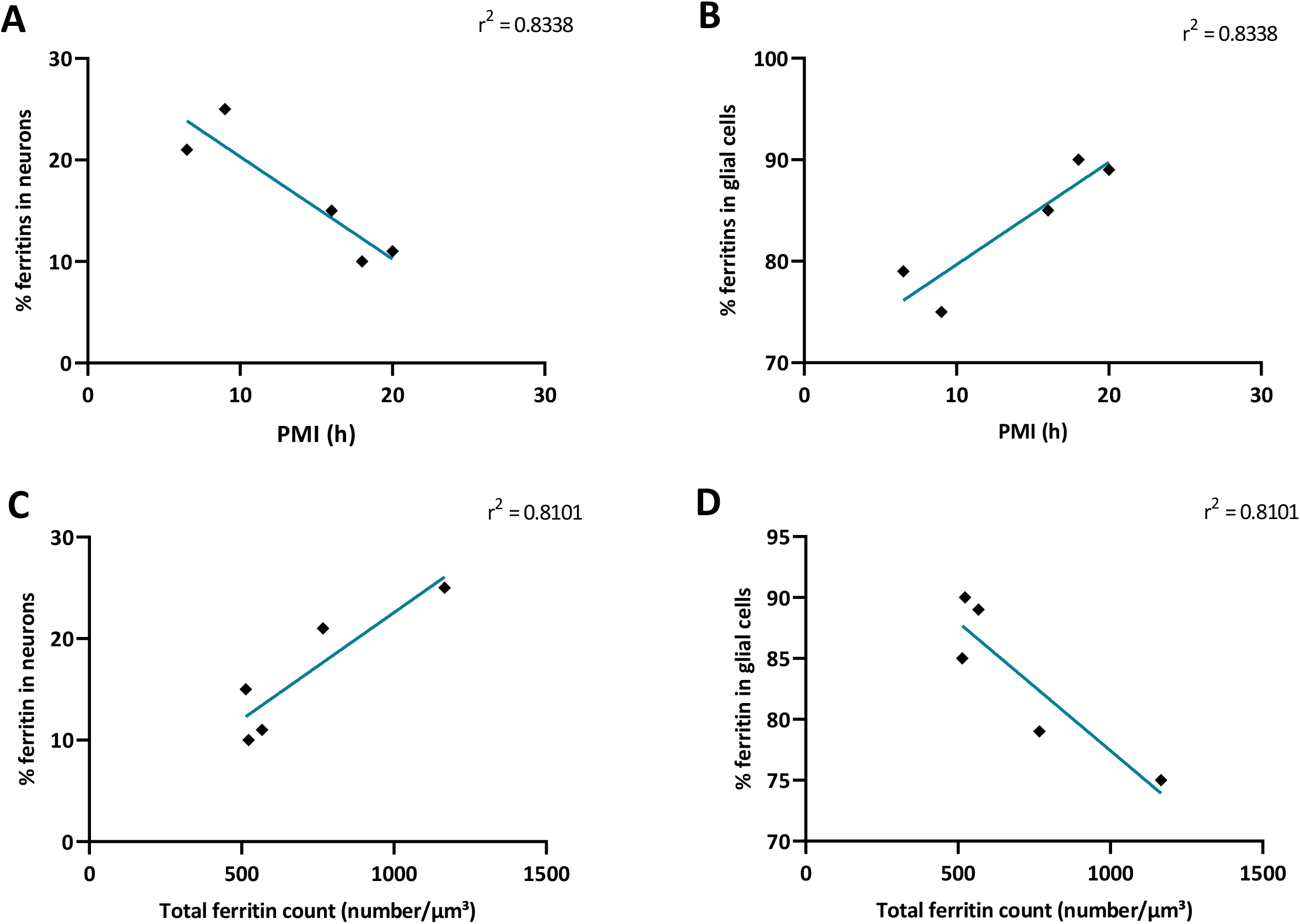
The percentage of ferritin in neurons varies with post-mortem time and with total ferritin concentration. **A**, Percentage of ferritins in neurons from patients 1 to 5 plotted against PMI. Mean values of the percentage of ferritin concentration in neurons (EFTEM) reduce over time with increasing PMI, r^2^ = 0.8338, p = 0.0303. **B**, Percentage of ferritins in glial cells from patients 1 to 5 plotted against PMI. Mean values of the percentage of ferritin concentration in glial cells (EFTEM) reduce over time with increasing PMI, r^2^ = 0.8338, p = 0.0303. **C**, Mean values of the percentage of ferritins found in neurons in all brain regions from patients 1 to 5 were plotted against total ferritin count. The probability of ferritins being found in neurons rises with the total ferritin concentration, r^2^ = 0.8101, p = 0.0374. **D**, the probability of ferritins being found in glial cells decreases with the total ferritin concentration, r^2^ = 0.8101, p = 0.0374. Each dot represents the mean percentage of ferritin found in neurons in all brain regions measured in patients 1-5, respectively (n = 5). * p < 0.05, Linear correlation analysis.

To evaluate whether the cell-specific accumulation of ferritin is dependent on the total ferritin amount, we plotted the percentage of ferritin found in neurons against the total ferritin count in each sample. We observed that the percentage of ferritins stored in neurons was proportional to the total ferritin count. In other words, when the ferritin concentration increased in a patient, the probability of ferritins accumulating in neurons increased. This result indicates that, although the majority of ferritins was situated in glial cells, total ferritin load scaled with the proportion of ferritins found in neurons (Fig. 10C) and vice-versa for glial cells (Fig. 10D).

## Discussion

Our studies implemented analytical electron microscopy to determine the concentration of ferritin cores filled with iron and used ICP-MS and R_2_* relaxation rate from qMRI to determine the absolute iron concentration in the human brain. Samples from four brain regions of six deceased subjects were examined. The subjects had not suffered from a known neurological disorder and the samples were taken during routine autopsies.

Due to the electron density of the ferritins’ iron cores, they are directly visible in classical electron micrographs (reviewed in (Iancu, 2011)), but they can become obscured by the contrasting agent in routine contrasted sections (Iancu, 2011). Accordingly, we chose an energy filter to create iron maps for trustworthy ferritin core identification. The size (Iancu, 2011) and detectability of these clusters indicate that they correspond to the iron-loaded cores of ferritins. After identifying the ferritin cores, we compared the elemental maps with a classical electron micrograph of the same area for cell-type identification and ferritin localization. We were able to demonstrate that the putamen and the GP store more ferritin iron clusters than the FGM and the FWM, confirming a trend also seen in data obtained with both qMRI and ICP-MS.

The EFTEM method is based on visualizing iron clusters inside ferritin using analytical electron microscopy. Ferritins are known to vary in their iron load (Harrison and Arosio, 1996; Iancu, 2011; Jian et al., 2016) and those ferritin proteins that contain a low amount of iron remain undetectable using EFTEM. Instead of using a fixed detection threshold, this study relied on four independent researchers agreeing on the ferritin count from both iron L – elemental maps and iron L – jump-ratios. There is high variability in the data obtained using the EFTEM method due to the small sampling size. Hence, the mean concentrations of ferritin both from each patient and from each region were used for this study.

We were able to demonstrate that the putamen and GP had a higher mean concentration of detectable (iron-loaded) ferritins than the FGM and FWM. The basal ganglia as the storage site for iron have long been known (Hallgren and Sourander, 1958) and have since been confirmed in several different studies that used a variety of different methods (Maeda et al., 1997; Pfefferbaum et al., 2009; Krebs et al., 2014; Ramos et al., 2014). To validate our findings, we measured the iron concentration in adjacent pieces of tissue taken from the same patients with ICP-MS, and measured the proton relaxation rate R_2_*, using qMRI in the contralateral region of the same patients. Our study on iron concentration validates and confirms earlier studies in that the basal ganglia have both higher iron concentrations and a higher R_2_* relaxation rate than the frontal gray and white matter (Langkammer et al., 2010).

Our study found a high correlation between the R_2_* relaxation rate (qMRI) and the absolute iron concentration (ICP-MS), confirming an earlier, comparative study of both methods [16, 25]. In contrast, the mean ferritin concentration of all the brain areas did not correlate with the mean iron concentration, indicating that the number of iron-loaded ferritins was not directly dependent on the total iron concentration in each patient. It has to be borne in mind that the EFTEM method measures only those ferritin cores that are filled with a minimum threshold amount of iron (cargo), whereas both qMRI and ICMPS measure the total elemental iron load of the brain sample. We thus investigated the factors influencing the iron cargo load of ferritins and found that this load is highly dependent on the post-mortem interval: the longer the PMI, the fewer the iron-loaded ferritin cores found in a sample. This indicates that autolysis progresses very rapidly during the post-mortem time and leads to the unloading of iron from the ferritins’ core. Two factors may contribute to unloading: either the ferritin shell shielding the iron core is damaged during autolysis causing the leakage of its cargo, or the ferritin holoprotein is decayed dissipating the iron. Unloading of ferritin cores explains why the absolute iron concentrations remained independent of the post-mortem time, whereas the number of iron-loaded ferritin cores decreased with increasing post-mortem time. We thus can only speculate as to the cellular mechanisms that govern the dispersion or decaying of ferritin during the post-mortem interval. Further studies will be necessary to detect possible changes in the oxidation state during the unloading or decay of ferritins.

Our second aim was to reveal the role of different cell types in preserving ferritin during autolysis. We were able to determine the cell type (neuron, oligodendrocyte, astrocyte, or other glial cells) that contained most (86 %) of the ferritins we had detected in five out of the six patients. We found that most of the iron in the human brain is stored in oligodendrocytes, but iron is also found in other or unidentifiable glial cells and neurons. Owing to the fact that oligodendrocytes are the myelinating cells, it is apparent that more iron is stored in oligodendrocytes since it is used as a co-factor in lipid biosynthesis (Connor and Menzies, 1996; Todorich et al., 2009). Earlier studies support our results using a variety of different techniques (Connor et al., 1990; Quintana et al., 2004; Quintana et al., 2006; Meguro et al., 2008).

In examining the cell types that are involved in iron-filled ferritin loss during autolysis, we found that the loss occurs at a higher rate in neurons than in glial cells, as schematically depicted in Fig. 11. Interestingly, we found that the probability of the ferritins being stored in neurons rather than glial cells scales positively with total ferritin concentration in each patient. To our knowledge, this is the first study that shows that the iron–filled ferritin increases faster in neurons than in glial cells with increasing total ferritin concentration.

**Figure 11.**
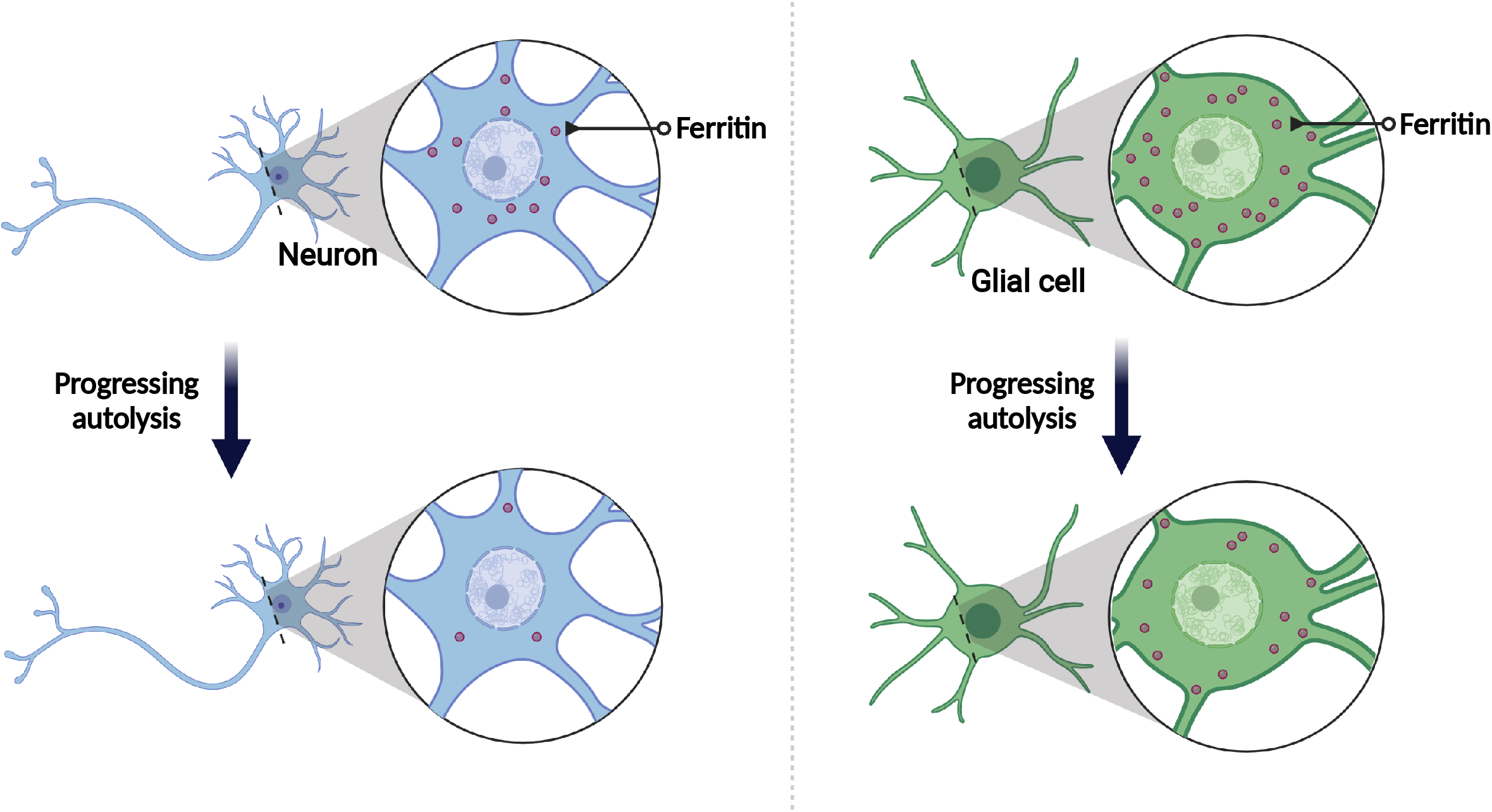
The rate of loss of the iron-filled ferritin cores during autolysis is higher in neurons than in glial cells.

Neurons were shown to have a higher ferritin H isoform distribution compared with glial cells(Han et al., 2002; Muñoz et al., 2011). Previous studies have shown that iron mediates the upregulation of ferritin H isoform in neurons(Nahirney and Tremblay, 2021). Thus, we speculate that iron overload might lead to increased production of ferritin H in neurons. This could also be a potential explanation for enhanced ferritin load increase in neurons when a higher concentration of iron is present in the brain.

In summary, we found high involvement of neurons in the storage of ferritins. Moreover, iron-loaded ferritins are lost post-mortem, in a yet unknown process, and loss of iron from ferritin cores occurs at a higher rate in neurons than in glial cells.

## Footnotes

- This work was funded by the Austrian Science Foundation [FWF], grant P 29370, and approved by the Ethics Committee of the Medical University of Graz, votum number 28-549 ex 15/16.
- We are grateful to Dr. Sigurd Lax, for advice and for contributing material, Dr. Peter Simmons (Newcastle University, UK), for proofreading and language corrections, Mariella Sele, for processing the samples and for initially evaluating most of the samples, Elisabeth Bock, Elisabeth Pritz, Luca Schmid, Susanne Sumerauer and Nina Schlögl for expert technical assistance. We especially thank Manuel Hündler for ferritin isolation.
- Figures 1 and 11 are created with BioRender.com
- The authors declare no competing financial interests.
- Correspondence should be addressed to Snježana Radulović at

## Abbreviations

EFTEM: energy-filtered transmission electron microscopy
EM: electron microscopy
FGM: frontal gray matter
FWM: frontal white matter
GP: globus pallidus
ICP-MS: inductively coupled plasma mass spectrometry
MR: magnetic resonance
PMI: *post-mortem* interval
qMRI: quantitative magnetic resonance imaging

## Author contributions

SS, SnR, and GL planned and performed the experiments at the electron microscope, and computed elemental maps and jump ratios. SS, SnR, GL, and SL analyzed and quantified the raw data from elemental maps. JH, AMTB provided and assessed the brain tissue. StR, SE, and CB performed all qMRI experiments and analysis. WG performed ICP-MS experiments and analysis. PS proofread the manuscript. GL designed the project and study plan. All authors revised, read, and approved the manuscript.

## References

Bilgic B, Pfefferbaum A, Rohlfing T, Sullivan EV, Adalsteinsson E (2012) MRI estimates of brain iron concentration in normal aging using quantitative susceptibility mapping. Neuroimage 59:2625–2635.

Codazzi F, Pelizzoni I, Zacchetti D, Grohovaz F (2015) Iron entry in neurons and astrocytes: a link with synaptic activity. Front Mol Neurosci 8:18.

Connor JR, Menzies SL (1996) Relationship of iron to oligondendrocytes and myelination. Glia 17:83–93.

Connor JR, Menzies SL, St Martin SM, Mufson EJ (1990) Cellular distribution of transferrin, ferritin, and iron in normal and aged human brains. J Neurosci Res 27:595–611.

(2016) ESMRMB 2016, 33rd Annual Scientific Meeting, Vienna, AT, September 29 - October 1: ePoster / Paper Poster / Clinical Review Poster / Software Exhibits. MAGMA 29 Suppl 1:401–475.

Finazzi D, Arosio P (2014) Biology of ferritin in mammals: an update on iron storage, oxidative damage and neurodegeneration. Arch Toxicol 88:1787–1802.

Friedman A, Arosio P, Finazzi D, Koziorowski D, Galazka-Friedman J (2011) Ferritin as an important player in neurodegeneration. Parkinsonism Relat Disord 17:423–430.

Giometto B, Gallo P, Tavolato B (1993) Transferrin Receptors in the Central Nervous System. In: Methods in neurosciences (Conn PM, ed), pp 122–134. Elsevier.

Hallgren B, Sourander P (1958) The effect of age on the non-haemin iron in the human brain. J Neurochem 3:41–51.

Halliwell B (2006) Oxidative stress and neurodegeneration: where are we now? J Neurochem 97:1634–1658.

Han J, Day JR, Connor JR, Beard JL (2002) H and L ferritin subunit mRNA expression differs in brains of control and iron-deficient rats. J Nutr 132:2769–2774.

Harrison PM, Arosio P (1996) The ferritins: molecular properties, iron storage function and cellular regulation. Biochimica et Biophysica Acta (BBA) - Bioenergetics 1275:161–203.

Iancu TC (2011) Ultrastructural aspects of iron storage, transport and metabolism. J Neural Transm (Vienna) 118:329–335.

Jian N, Dowle M, Horniblow RD, Tselepis C, Palmer RE (2016) Morphology of the ferritin iron core by aberration corrected scanning transmission electron microscopy. Nanotechnology 27:46LT02.

Jiang X, Stockwell BR, Conrad M (2021) Ferroptosis: mechanisms, biology and role in disease. Nat Rev Mol Cell Biol 22:266–282.

JoVE Video Dataset.

Krebs N, Langkammer C, Goessler W, Ropele S, Fazekas F, Yen K, Scheurer E (2014) Assessment of trace elements in human brain using inductively coupled plasma mass spectrometry. J Trace Elem Med Biol 28:1–7.

Langkammer C, Krebs N, Goessler W, Scheurer E, Ebner F, Yen K, Fazekas F, Ropele S (2010) Quantitative MR imaging of brain iron: a postmortem validation study. Radiology 257:455–462.

Lewis AJ, Genoud C, Pont M, van de Berg WD, Frank S, Stahlberg H, Shahmoradian SH, Al-Amoudi A (2019) Imaging of post-mortem human brain tissue using electron and X-ray microscopy. Curr Opin Struct Biol 58:138–148.

Maeda H, Sato M, Yoshikawa A, Kimura M, Sonomura T, Terada M, Kishi K (1997) Brain MR imaging in patients with hepatic cirrhosis: relationship between high intensity signal in basal ganglia on T1-weighted images and elemental concentrations in brain. Neuroradiology 39:546–550.

Marengo-Rowe AJ (2006) Structure-function relations of human hemoglobins. Proc (Bayl Univ Med Cent) 19:239–245.

Meguro R, Asano Y, Odagiri S, Li C, Shoumura K (2008) Cellular and subcellular localizations of nonheme ferric and ferrous iron in the rat brain: a light and electron microscopic study by the perfusion-Perls and -Turnbull methods. Arch Histol Cytol 71:205–222.

Muñoz P, Humeres A, Elgueta C, Kirkwood A, Hidalgo C, Núñez MT (2011) Iron mediates N-methyl-D-aspartate receptor-dependent stimulation of calcium-induced pathways and hippocampal synaptic plasticity. J Biol Chem 286:13382–13392.

Nahirney PC, Tremblay M-E (2021) Brain Ultrastructure: Putting the Pieces Together. Front Cell Dev Biol 9:629503.

Pfefferbaum A, Adalsteinsson E, Rohlfing T, Sullivan EV (2009) MRI estimates of brain iron concentration in normal aging: comparison of field-dependent (FDRI) and phase (SWI) methods. Neuroimage 47:493–500.

Quintana C, Bellefqih S, Laval JY, Guerquin-Kern JL, Wu TD, Avila J, Ferrer I, Arranz R, Patiño C (2006) Study of the localization of iron, ferritin, and hemosiderin in Alzheimer’s disease hippocampus by analytical microscopy at the subcellular level. J Struct Biol 153:42–54.

Quintana C, Cowley JM, Marhic C (2004) Electron nanodiffraction and high-resolution electron microscopy studies of the structure and composition of physiological and pathological ferritin. J Struct Biol 147:166–178.

Ramos P, Santos A, Pinto NR, Mendes R, Magalhães T, Almeida A (2014) Iron levels in the human brain: a post-mortem study of anatomical region differences and age-related changes. J Trace Elem Med Biol 28:13–17.

Sele M, Wernitznig S, Lipovšek S, Radulović S, Haybaeck J, Birkl-Toeglhofer AM, Wodlej C, Kleinegger F, Sygulla S, Leoni M, Ropele S, Leitinger G (2019) Optimization of ultrastructural preservation of human brain for transmission electron microscopy after long post-mortem intervals. Acta Neuropathol Commun 7:144.

Todorich B, Pasquini JM, Garcia CI, Paez PM, Connor JR (2009) Oligodendrocytes and myelination: the role of iron. Glia 57:467–478.

Wagner KR, Sharp FR, Ardizzone TD, Lu A, Clark JF (2003) Heme and iron metabolism: role in cerebral hemorrhage. J Cereb Blood Flow Metab 23:629–652.

Wang J, Pantopoulos K (2011) Regulation of cellular iron metabolism. Biochem J 434:365–381.

Ward RJ, Zucca FA, Duyn JH, Crichton RR, Zecca L (2014) The role of iron in brain ageing and neurodegenerative disorders. The Lancet Neurology 13:1045–1060.

Wernitznig S, Reichmann F, Sele M, Birkl C, Haybäck J, Kleinegger F, Birkl-Töglhofer A, Krassnig S, Wodlej C, Holzer P, Kummer D, Bock E, Leitinger G (2019) An Unbiased Approach of Sampling TEM Sections in Neuroscience. J Vis Exp. doi:10.3791/58745.

